# Entropy of the resting state cortex in epilepsy

**DOI:** 10.1101/2025.10.02.680016

**Authors:** Kirandeep Kaur, Jack O’Brien-Cairney, Gaurav Singh, Manoj Upadhya, Aphrodite Chakraborty, Sarat P Chandra, Harald Prüss, Hans-Christian Kornau, Dietmar Schmitz, Gavin Woodhall, Richard Rosch, Maia Angelova, Stefano Seri, Manjari Tripathi, Sukhvir K Wright, Caroline Witton

## Abstract

**Background:** Epilepsy has long been conceptualised as a disorder in which aberrant brain dynamics extend beyond the epileptogenic zone. Evidence demonstrates that loss of entropy is a generic feature of pathological dynamics in the brain, including the ictal state. However, the impact of recurrent seizures on entropy in the interictal state remains unknown.

**Methods:** Resting state magnetoencephalography (MEG) scans and resection masks of 32 individuals with epilepsy who had Engel I outcome post-surgery were retrospectively retrieved. Using co-registered FreeSurfer parcellations, we reconstructed the source localised MEG time series and computed sample entropy for 114 regions of interest. We then tested the association of entropy with the resected volume of the brain, and additional clinical variables including the age of seizure onset, seizure frequency and duration of epilepsy. To further understand the temporal relationship between seizure onset and entropy in the interictal state, we collected and computed sample entropy for week-long EEG traces from leucine-rich glioma inactivated 1 monoclonal antibody (LGI1-mAb) rodent models of autoimmune encephalitis (n=5) and control rats (n=5).

**Results:** In individuals with epilepsy, a lower age of seizure onset was associated with lower mean sample entropy of the whole cortex (Spearman’s rho =0.60, p<0.001; partial correlation =0.41, p=0.021). Entropy did not differ between the resected and non-resected regions of the brain. Furthermore, LGI1-mAb treated rodents showed a persistent decrease in sample entropy as compared to control rats, after the onset of seizures, and this difference was greatest during periods of highest seizure frequency (p<0.001).

**Conclusion:** Recurrent seizures are associated with a persistent decrease in entropy, even in the interictal state, and this decrease was found to be most profound and affecting the whole cortex in patients who had a lower age of seizure onset.

## Introduction

In nature, complex systems are constituted of a large number of interacting parts which collectively give rise to patterns of functioning and/or behaviour that cannot be explained by the laws governing the constituent parts themselves (Estrada 2023; Ladyman, Lambert, and Wiesner 2013). For example, starling murmuration (an observable phenomenon where thousands of starling birds fly together in synchronized and continuously shifting patterns), cannot be explained by the behaviour of a single starling alone (King and Sumpter 2012). Likewise, the human brain is another example of a complex system where individual neurons collectively give rise to higher order functions like language and cognition(Baars and Gage 2010; Gallup Jr. 1982).

Many complex systems including the mammalian brain are hypothesized to function in a state of *criticality*, which is an intermediate state between complete order and randomness (Plenz et al. 2021; Rickles, Hawe, and Shiell 2007) and where complexity is maximal(Huberman and Hogg 1986). It is posited that systems in a state of criticality have the greatest capacity for adaptation and information processing(Hesse and Gross 2014; Rohlf and Bornholdt 2009). However, loss of complexity in neuronal activity is a hallmark of natural ageing, altered states of consciousness as well as selected brain disorders (Dauwels et al. 2011; Keshmiri 2020; Manor and Lipsitz 2013; Yang et al. 2015).

Complexity of a system is challenging to quantify (Clark and Jacques 2012). Entropy, which is fundamentally regarded as the degree of order or randomness is a system has been used to measure the complexity in natural systems (S. M. Pincus 1995:1). At a foundational level, complexity is considered to be an emergent property of the continuously increasing entropy in the universe (Modis 2022). There exist several interpretations of entropy for neural systems ranging from the degree of disorder in neural signals, to the information processing capacity of the brain(Balibrea 2016; Cieri et al. 2021; Keshmiri 2020). However, entropy is most accurately interpreted as the number of varied states or configurations in which the brain as a neural network can exist(Boltzmann 2022; Saxe, Calderone, and Morales 2018; Zagha and McCormick 2014).

In patients with epilepsy, brain entropy falls sharply during an ictal event because neural activity during seizures becomes more synchronous and predictable (Mammone et al. 2015; Nicolaou and Georgiou 2012; Tibdewal et al. 2017). Evidence from animal models of epilepsy suggests that recurrent seizures can lead to temporary or even permanent functional changes in synaptic activity by changing its threshold(Hempel et al. 2000; Majak and Pitka nen 2004; Yang et al. 2021). Therefore, we hypothesized that given its propensity for recurring seizures, the entropy of the epileptogenic zone would be lower as compared to the non-epileptogenic brain regions. The regional differentiation may serve as an additional tool to localize the epileptogenic zone.

To test this hypothesis, we computed sample entropy from source localized resting state (interictal) magnetoencephalography (MEG) data of patients who had undergone epilepsy surgery and had Engel I outcome for at least 3 years post-surgery. Sample entropy is a robust measure of the degree of irregularity or complexity in a time-series signal (Delgado-Bonal and Marshak 2019; Namdari and Li 2019). In addition, entropy derived from human data was tested for its association with other clinical variables such as age of seizure onset and duration of epilepsy. To further understand the causal link between seizures and abnormal interictal brain dynamics, we evaluated how entropy changes with the onset of epileptogenesis by analysing electroencephalography (EEG) traces from leucine-rich glioma inactivated 1 monoclonal antibody (LGI1-mAb) rodent models of autoimmune encephalitis and control rats. LGI1-mAb rodents are an established model of autoimmune epilepsy (Upadhya et al. 2025), and in this study they were used to assess any change in entropy, before and after the onset of seizures.

## Methods

### Acquisition and pre-processing of MEG data

All clinical data was collected retrospectively with approval from the institutional ethics committee of All India Institute of Medical Sciences, New Delhi, India. The clinical history, and MEG data were retrieved for 36 patients who had Engel I outcome (complete seizure freedom) for a duration of at least 3 years following surgery. Retrieving data only from patients who had complete seizure freedom post-surgery reduces uncertainty regarding the source of epileptogenic activity.

The source-localized MEG data, parcellated structural MRI, and the resection masks used here were reanalysed from a previously reported cohort (Kaur et al. 2023), and included 32 retrievable datasets. An overview of the methodology for MEG source analysis and creation of resection mask is given below.

T1-weighted MRIs were pre-processed in FreeSurfer (Fischl 2012), using the Lausanne parcellation scheme (Hagmann et al. 2008). For this study, we conducted our analysis over 114 neocortical regions of interest (ROIs), excluding any deep brain structures such as hippocampus, thalamus and amygdala. The following steps were used to create masks for the regions which were resected during surgery: (i) FSL FLIRT was used to linearly co-register the postoperative MRI/CT (displaying the resection cavity) and the preoperative Freesurfer parcellated MRI (Jenkinson et al. 2002; Jenkinson and Smith 2001). In cases where only the post-operative CT was available, the ITK-SNAP tool was used to adjust the contrast (Yushkevich et al. 2006), (ii) masks were manually drawn over the resection cavity in FSLView and smoothened using the *fslmaths* toolbox.

Resting-state MEG was acquired for all patients preoperatively in a supine position, using a 306-channel Elekta Neuromag system (Helsinki). MEG was recorded at 1KHz sampling frequency and corrected for head movement artefacts using the method of temporal Signal Space Separation (tSSS)(Taulu and Simola 2006). Further details of MEG acquisition protocols for these patients have been described previously (Kaur et al. 2021; Tripathi et al. 2021). Raw MEG data were down-sampled to 500Hz for computational efficiency. All further pre-processing was done in Brainstorm software (Tadel et al. 2011). The MEG data were co-registered with the parcellated structural MRI and band-pass filtered between 1-100Hz. It was further notch-filtered at 50Hz using a second-order IIR (infinite impulse response) filter. Signal Space Projection (SSP) was used to remove eye movement and electrocardiographic artefacts (Tesche et al. 1995; Uusitalo and Ilmoniemi 1997). The next step was to generate a single MEG time series for each ROI across the whole cortex. This was done by constructing a head model from the structural MRI for every patient using the overlapping spheres method. The sLORETA algorithm (Pascual-Marqui 2002) was used to create a source model of the MEG signal across 114 neocortical regions.

### Patient Characteristics

The clinical and demographic characteristics of 32 patients are provided in Table I. As mentioned previously, all patients had reported an Engel I outcome for at least 3 years post-surgery.

**Table I:** Clinical and demographic characteristics of patients.

| Clinical and demographic characteristics |  |
| --- | --- |
| Age [Mean ± SD] | 15.91 ± 8.86 |
| Duration of epilepsy in years [Mean ± SD] | 8.09 ± 6.35 |
| Age at seizure onset in years [Median(IQR)] | 5.75 (1.5-13.0) |
| Seizure frequency/ week [Median (IQR)] | 7.0 (1.44 - 19.25) |
| Sex<br>Female (%)<br>Male (%) | 12 (37.5)<br>20 (62.5) |
| Lateralisation of the region resected during surgery<br>Left (%)<br>Right (%) | 14 (43.75)<br>18 (56.25) |

### Acquisition and pre-processing of rodent EEG

Wistar rats, at postnatal age day 21 (P21) were used to gather electroencephalogram (EEG) data and recorded for a period of 21 days (N=10). The rats were infused with patient-derived anti-LGI1 monoclonal antibodies or control antibodies (non-brain reactive) intracerebroventricularly via two subcutaneously implanted osmotic pumps (Alzet model 1007D, volume 100uL, flow rate 0.5uL per hour, duration 7 days). Out of the 10 rats used, five were infused with control antibodies and five were infused with human derived monoclonal anti-LGI1 antibodies (Upadhya et al. 2025). Paired video-EEG recordings were collected using a previously standardised protocol (Kreye et al. 2021; Wright et al. 2021).

To construct an appropriate analogue to human interictal MEG data, ictal events within the rodent EEG were first identified using a standardized protocol (Upadhya et al. 2024, 2025) and omitted from further analysis.

### Computing Entropy

We used sample entropy (SE) to measure the entropy of MEG source localized time-series and rodent EEG. Sample entropy measures the degree to which a time-series has similar or self-repeating patterns of a given length (Richman and Moorman 2000). Mathematically, sample entropy is an estimate of the conditional probability that two sequences in a time-series signal, which are similar for ‘m’ points, remain similar (within tolerance) after the addition of the next point ‘n’, excluding any self-counting (i.e. matching a sequence to itself) (Delgado-Bonal and Marshak 2019; Richman and Moorman 2000). The lowest possible value of sample entropy is 0 (indicating a higher degree of regularity or self-repetition in a time-series), while greater values indicate greater irregularity or complexity. Conceptually, a lower value of entropy in a neural signal would indicate that the underlying neurons are accessing similar states repeatedly over the measured time course.

SE is very similar in computation to another measure of regularity of a time-series, approximate entropy, except that it excludes self-matching of sequences (S. M. Pincus 1995a;

S. Pincus 1995b).This makes sample entropy a more robust tool for measuring the presence of stochasticity (randomness) or regularity in a time-series over shorter durations (Richman and Moorman 2000).

SE was computed for the first 9 minutes (540s) of each of the 114 ROI time series per patient. This duration was concluded to be sufficient since the mean sample entropy across patients was found to stabilize and remain approximately constant after 360s (6 minutes) (eFigure1). Given a sampling frequency of 500Hz, 9 minutes of data corresponded to 270,000 data points per ROI. In line with recommendations by Bonal *et al*.(Delgado-Bonal and Marshak 2019), the tolerance, ‘r’, for accepting matches, i.e., the degree to which a sequence can deviate from another and still be considered similar, was set as 0.2 SD (20% of the standard deviation of the time series). The length of the sequences (*m*) to be compared, was set as three. Entropy was computed from each neural time series signal in MATLAB 2021a using the function developed by Lord et al (Lord, 2024).

The rodent EEG on the other hand was epoched into hour-long recordings, and the sample entropy was computed for each of these epochs using the same parameters and function. The total duration of rodent EEG recordings totalled 192 hours (∼8 days).

### Statistical Analysis

Statistical analysis was conducted in R (version 2023.06.2) and MATLAB_R2021a.

For human MEG data, non-parametric tests were used to test the difference in average sample entropy between the hemispheres ipsilateral and contralateral to the site of resection, and the resected and non-resected regions of the brain. Multiple linear regression was used to test the association between sample entropy and seizure frequency, duration of epilepsy, age of seizure onset, sex, age at the time of MEG scan, lateralization of epileptogenic zone, lobe of resection and the presence of focal to bilateral tonic-clonic seizures. Since the number of predictor variables in this case was very high (n=11), stepwise regression in backward direction was used to find the clinical variables which best explained the variation in sample entropy. Backward stepwise regression begins with a model which has all predictor variables and iteratively removes the least statistically significant variable, one at a time till the best-fit model is achieved according to the Akaike information criterion (AIC). The best-fit model as per AIC is the one which explains the maximum possible variance in the dependent variable using the fewest independent variables. In cases where a confounding variable was found to impact both the independent and dependent variables, partial correlation was used to adjust for the confounding variable. This was done by regressing out the confounding variable from the dependent and independent variables, and then testing the residuals of these models for the presence of any significant correlations.

For rodents, Wilcoxon signed-rank test was used to determine the difference in average sample entropy between control and LGI1-mAb infused rats over a 24-hour period. The Tukey-Kramer procedure was used to adjust for multiple comparisons. Cross-correlation function was used to find the lagged correlation between the mean seizure frequency and sample entropy per hour. The ‘astsa’ package in R was used to detrend the time-series of seizure frequency and sample entropy (Stoffer 2012).

## Results

### Human MEG data

#### Entropy is highly variable between patients

The mean sample entropy of the whole cortex across patients was 0.84 ± 0.12. Interestingly, the variability of mean sample entropy between patients (SD=0.12) was significantly greater than the variability of the mean sample entropy between ROIs (SD=0.04). This indicates that the entropy of the whole cortex differed majorly from patient to patient, but did not show as much variability between different regions of the same cortex.

Therefore, contrary to our hypothesis, we did not find any significant difference in the mean sample entropy (SE) of the resected and non-resected regions of the brain. In addition, mean SE did not differ significantly between the hemisphere ipsilateral and contralateral to the site of resection.

#### Age of seizure onset is a predictor of entropy

To test the degree to which clinical characteristics of patients can predict their average SE across the cortex, we used stepwise regression analysis in the backward direction till the lowest possible AIC score was reached. We found that the best predictors of mean entropy across cortex were the age of seizure onset, lateralization of epileptogenic zone (hemisphere) and if at least one of the resected regions were in the parietal cortex. These variables together explained 47.33% of the variance in sample entropy (R2 = 0.4733; F(3,28)= 8.38, p = 0.0004) (Table 2).

**Table 2:** Multiple linear regression of mean sample entropy vs clinical characteristics.

|  |  |
| --- | --- |
| Dependent variable: |  |
| Mean sample entropy |  |
| Age of onset | 0.010***<br>(0.002) |
| Hemisphere (L/R) | 0.058<br>(0.032) |
| Parietal lobe resection | -0.101<br>(0.055) |
| Constant | 0.739***<br>(0.029) |
| Observations | 32 |
| R2 | 0.473 |
| Adjusted R2 | 0.417 |
| Residual Std. Error | 0.090 (df = 28) |
| F Statistic | 8.388*** (df = 3; 28) |

Note: \*\*p<0.05; \*\*\*p<0.01

#### The table shows the value of the intercept for each predictor variable and its standard error in brackets

The age of onset was the strongest predictor of SE (coefficient = 0.01, p <0.001), while the hemisphere of resection and resection in the parietal cortex did not individually reach the threshold of significance in the final regression model. Thus, we can conclude that SE increases by an average of 0.01 units for every year added to the age of epilepsy onset. A scatterplot between the age of onset and the average entropy across the cortex is shown in Figure 1A. In addition age of onset was found to be correlated not only with the variability of entropy between patients, but also between regions of the cortex (range of SE) within a patient (rho = −0.5, p =0.004)(Figure 1B). Duration of epilepsy was not significantly associated with the mean or the range of entropy. The association between entropy and the age of seizure onset remained significant even after adjusting for the patient’s age at the time of scan (rho = 0.41, p=0.21)(Figure 2). Duration of epilepsy was not significantly associated with the mean or the range of entropy.

**Figure 1.**
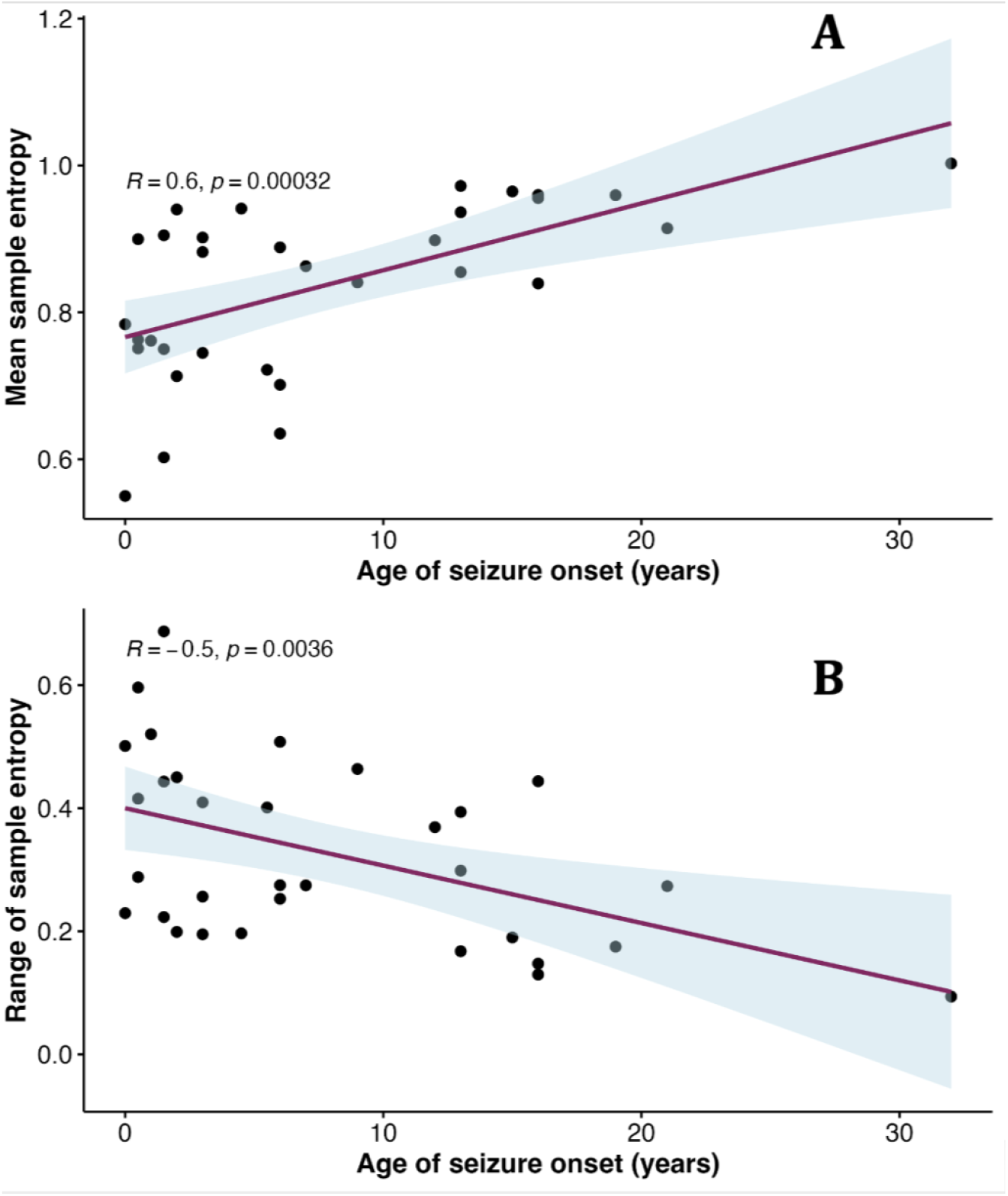
Age of seizure onset (years) correlates with the mean and range of sample entropy in the whole cortex. A) The scatter plot shows the association between average sample entropy across 114 regions of interest in each patient and their age of seizure onset in years. Each data point corresponds to one patient. The correlation coefficient (Spearman’s rho) was equal to 0.6 (p<0.001). The maroon line depicts the regression fit. B) Similarly the range (maximum-minimum) of entropy values within a patient was negatively correlated with their age of onset (Spearman’s rho = −0.49, p =0.005). This demonstrates that lower age of epilepsy onset is associated with greater variability of resting state sample entropy between different regions of the cortex.

**Figure 2.**
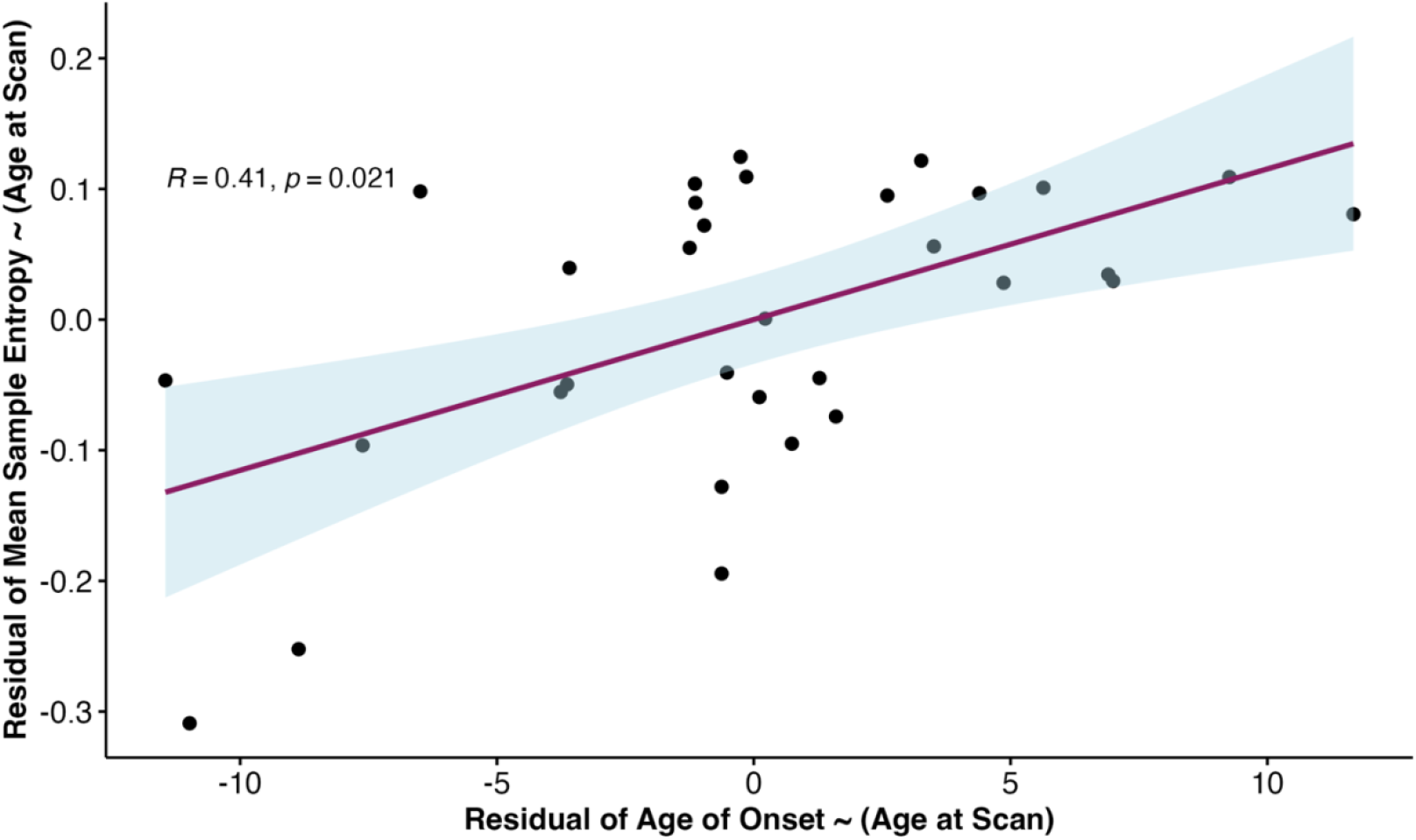
Partial correlation between age of seizure onset and mean sample entropy after adjusting for the age at scan. The scatter plot shows the correlation between the residuals of age of onset and the residuals of the mean sample entropy after a regressing out the age at the time of MEG scan. The correlation coefficient (Spearman’s rho) was equal to 0.41 and remained statistically significant (p=0.02).

Together, these results indicate that an earlier age of onset of seizures lowers the entropy of the whole cortex, and this alteration is sustained during the interictal state as well.

### Rodent Data

#### Sample Entropy of LGI1-mAb infused rodent EEG differs from controls

Next, we sought to examine how seizure onset alters interictal entropy. Week-long *in vivo* single-channel EEG recordings were collected and analysed from an independent rodent model of autoimmune encephalitis (LGI1-mAb, n=5) and control rats (n=5).

EEG traces during interictal periods among LGI1-mAb infused rats expressed significantly lower values of sample entropy than time-matched controls after day 1 of recording (Figure 3). The first seizure was recorded on day 1 itself in all LGI1-mAb rats.

**Figure 3.**
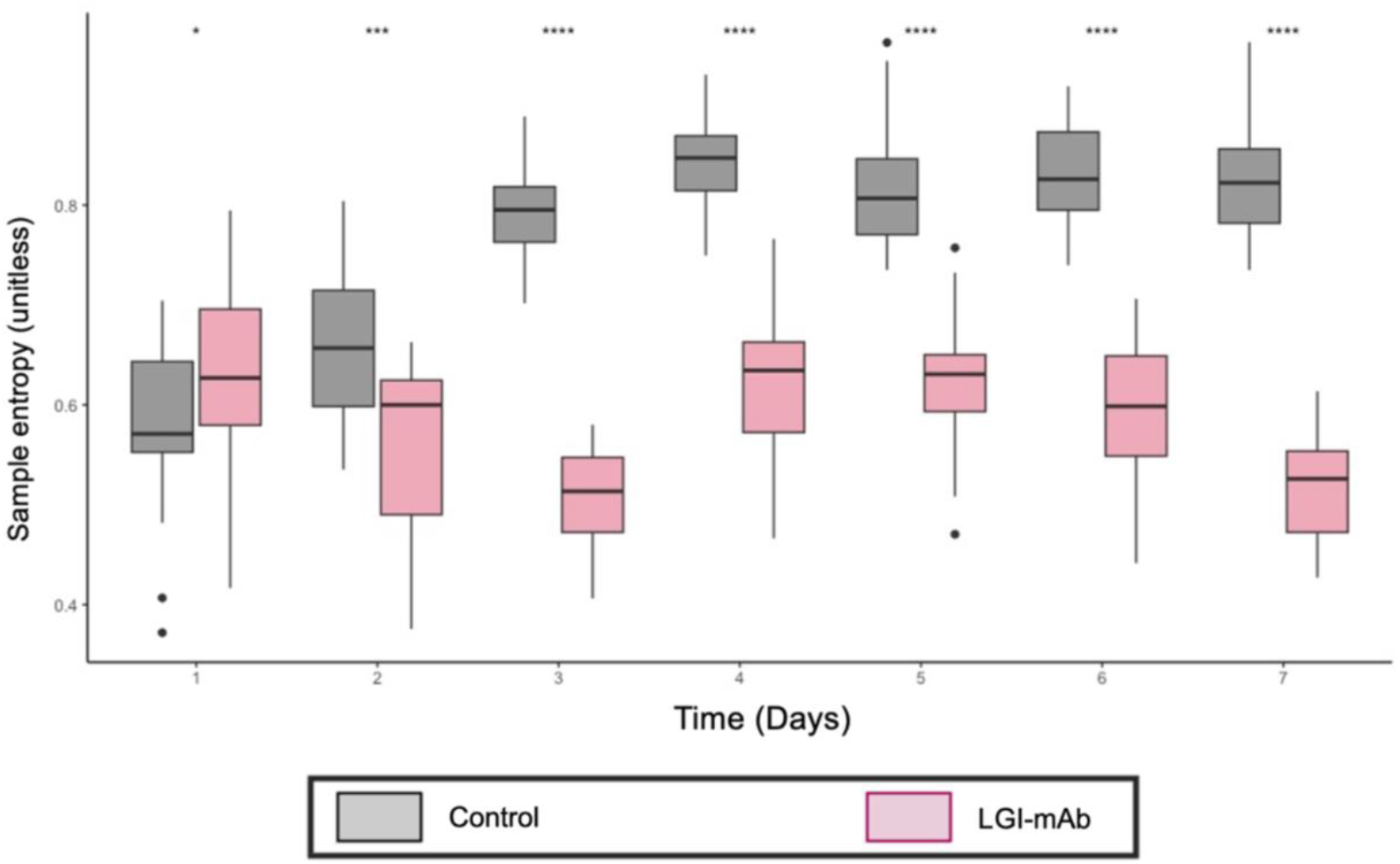
Difference between sample entropy of LGI1-mAb infused and control rats in interictal state. A box chart displaying the averaged, hourly sample entropy for each day in both rodent cohorts (N_Controls_ = 5, in grey; N_LGI1_ = 5 in pink). Each box and whisker plot represents the distribution of median sample entropy/hour in a 24-hour period, averaged across all rats in a particular cohort (i.e. 24 data points/day). The averaged sample entropy was significantly higher in the LGI1-mAb infused rats on day 1 of recording. However, from day 2 and onwards, the sample entropy of LGI1-mAb infused rats remained lower than that of controls. The first seizure was recorded on day 1 in all LGI1-mAb rodents. The statistical significance of the differences between disease cohorts and time-matched controls was tested using Wilcoxon signed-rank tests (^*^ :p<0.05; ^**^:p<0.01;^***^ :p<0.001; ^* * * *^ :p<0.0001).

### Seizure onset increases the entropy differences between LGI1-mAb infused and control rats

Next, we sought to understand how the entropy of LGI1-mAb infused and control rats evolves in time, specifically after the onset of seizures. We found that entropy in the interictal state does not differ significantly between LGI1-mAb and control rats before the onset of seizures (p=0.81) (Table 3). On the other hand, there was a sustained and significant decrease in SE of LGI1-mAb infused rats as compared to controls after the onset of seizures (Table3,Figure 3 & 4). Additionally, the difference in entropy shows a sustained trend even after cessation of the peak seizure frequency (Figure 4).

**Table 3:** Mean sample entropy in LGI1-mAb infused and control rats.

| LGI1-mAb infused rats (n=5) | Control rats (n=5) | p- value |
| --- | --- | --- |
| **For the entire duration of recording** |  |  |
| 0.58 ± 0.08 | 0.76 ± 0.11 | p<0.0001 |
| **Before seizure onset (First 6 hours, starting day 21)** |  |  |
| 0.52 ± 0.08 | 0.5 ± 0.1 | 0.81 |

| After seizure onset |  |  |
| --- | --- | --- |
| 0.58 ± 0.08 | 0.77 ± 0.1 | p<0.0001 |

**Figure 4.**
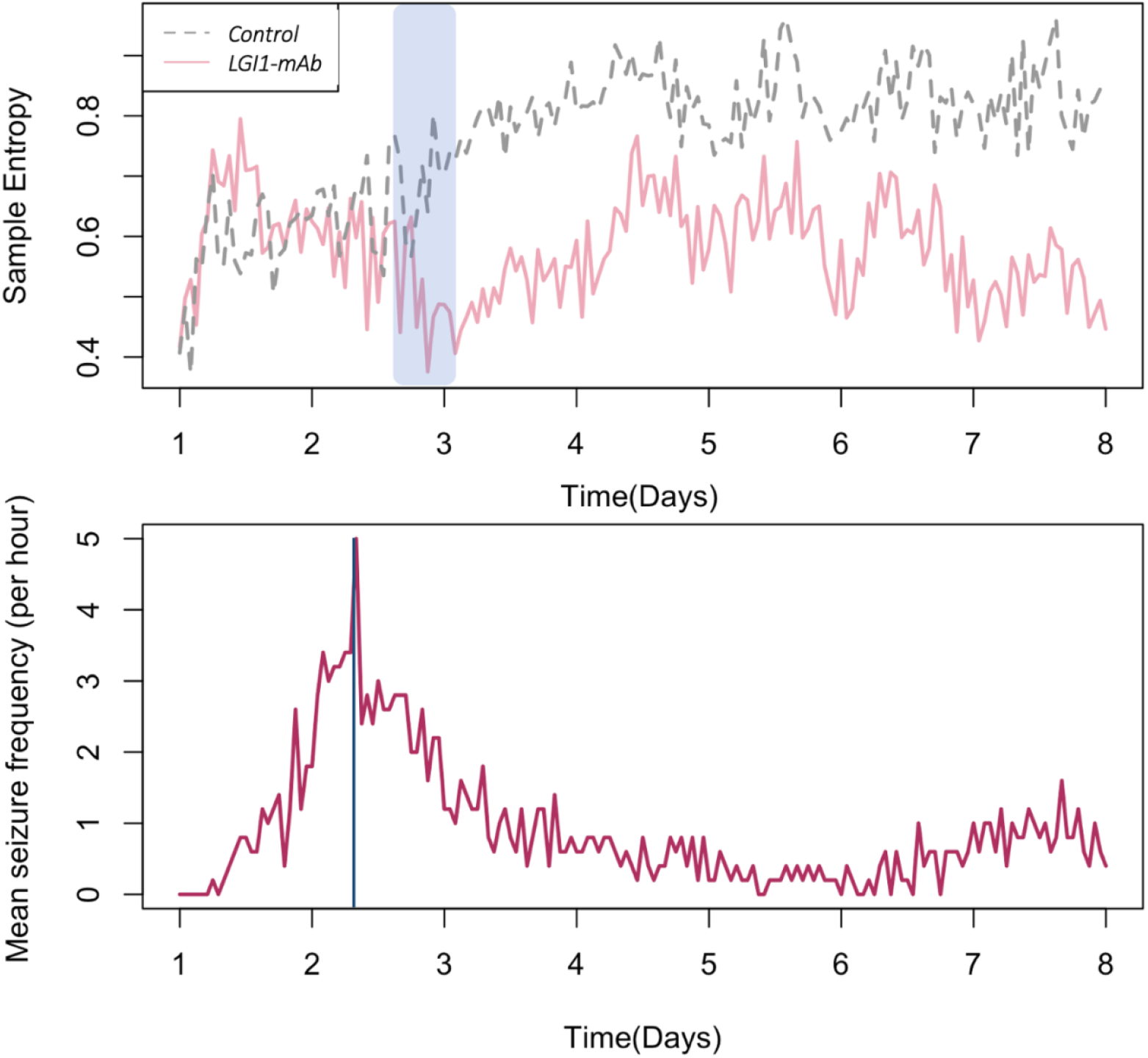
Time series plot of hourly median sample entropy (A) and mean seizure frequency/hour in LGI1-mAb rats(B) (A) Time series graph depicting the mean sample entropy per hour in LGI1-mAb infused rats (pink) and control rats (grey) (B) Time series graph of mean seizure frequency per hour in LGI1-mAb treated rats. We can see from the graph that a peak in seizure frequency between day 2 and 3 (blue solid line) is followed by a decrease in sample entropy (blue shaded region) in the LGI1-mAb infused rats.

Lastly, we used a cross-correlation function to test if there was a *lagged* correlation between seizure frequency and SE. Figure 5 shows the lagged correlation plot between the two time series. Cross-correlation (CCF) values tending to 1 or −1 demonstrate a strong positive or negative correlation between the two time series respectively, while values closer to 0 indicate a lack of correlation between the two variables. The plot indicates that there is a negative correlation between seizure frequency and sample entropy, and this relationship is strongest approximately at a lag of 9 hours. This is in line with the findings of Figure 4, wherein a peak in mean seizure frequency between day 2 and 3 (∼5 seizures/hour, 33 hours after treatment), is followed by the lowest observed value of mean SE (0.38) 13 hours later.

**Figure 5.**
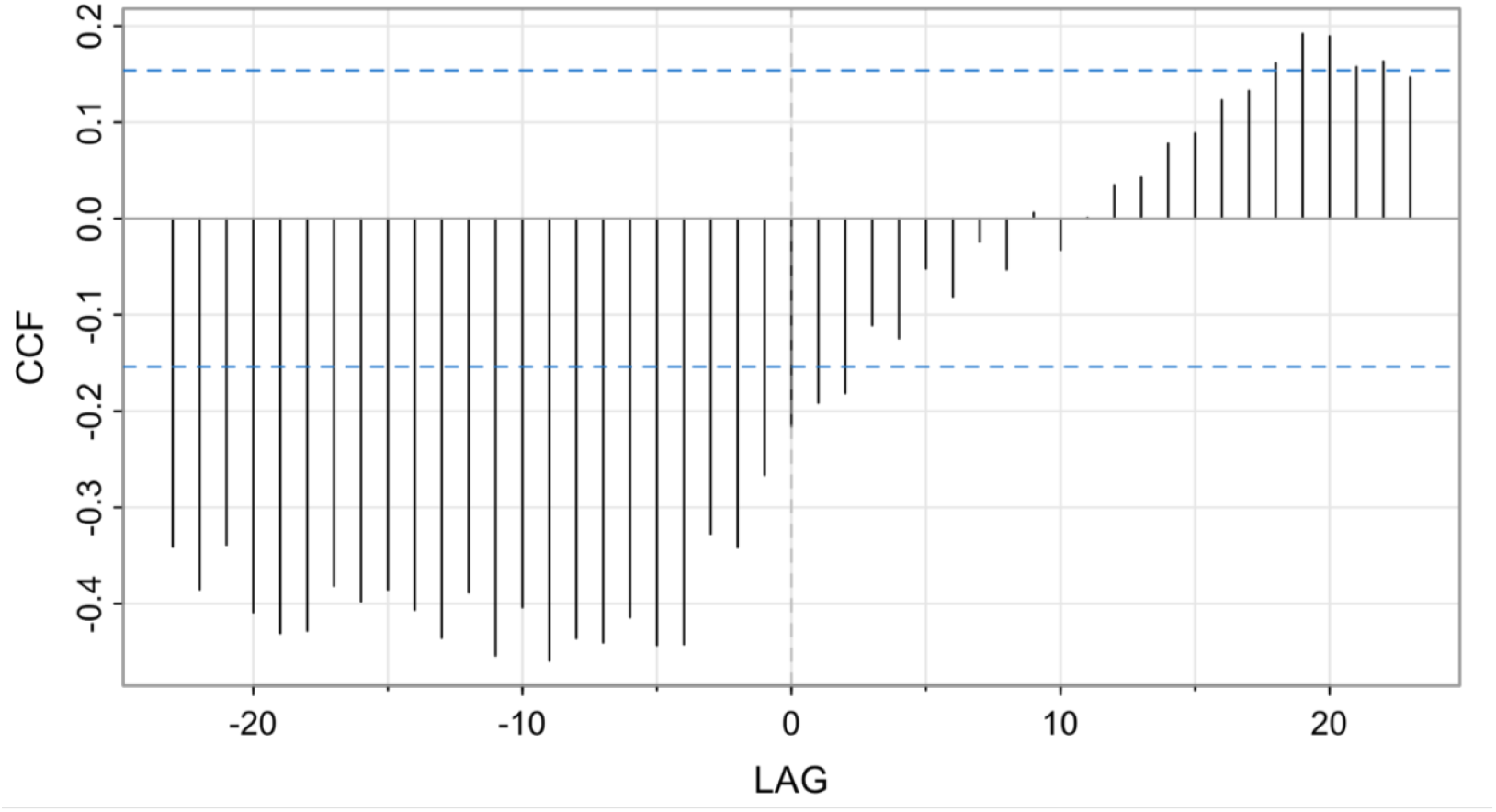
Lagged (cross-correlation) between mean seizure frequency/hour and mean sample entropy/hour. The x-axis shows the lag in hours between the two time series-(i) mean seizure frequency/ hour and (ii) averaged sample entropy/ hour in LGI1-mAb infused rodents (n=5). The y-axis shows the cross-correlation between the two series at different time lags. The dominant negative lag in this plot indicates that sample entropy ‘lags’ behind seizure frequency, or said another way-changes in seizure frequency may negatively affect sample entropy after a period of few hours. The strongest negative correlation (r = −0.46) was observed at a lag of 9 hours between seizure frequency and sample entropy time series.

## Discussion

In this study we explored the relationship between the onset of seizures and the levels of entropy in the resting state cortex of humans and rodents. While human MEG provides a spatially rich insight into the association between the history of ictal events and entropy at a single point in time, EEG data from the LGI1-mAb infused rodent model gives temporal information about how seizures may impact brain entropy.

In disagreement with our primary hypothesis, we did not observe any significant difference between the epileptogenic and non-epileptogenic regions of the brain with respect to SE. On the contrary, we found that SE showed little variability between different regions of the same cortex. These results are in contrast to a recent study which found that resection of interictal ‘low entropy zones’ derived from intracranial recordings was predictive of better Engel outcomes (degree of seizure remission post-surgery)(N et al. 2025). However, from the human MEG recordings we found that the age of seizure onset is strongly correlated with, and predictive of, the mean sample entropy of the whole cortex. Key clinical and demographic characteristics like seizure frequency, duration of epilepsy, age at MEG scan and sex did not significantly explain the variance in the entropy of patients. Instead, a stepwise regression model showed that a lower age of onset apparently ‘locks’ the whole cortex into a state of lower entropy and greater variability of the same from one region to another. These results strengthen the hypothesis that epilepsy is a ‘network’ disorder, rather than a regional disorder of the brain (Scharfman et al. 2018). In line with these results, we found significantly lower sample entropy in the rodent model of LGI1 autoimmune encephalitis as compared to control rats on days with the highest recorded seizure activity. Moreover, this lowering of entropy in LGI1-mAb rats persisted even after the initial peak of seizure frequency had passed.

Entropy falls drastically during an ictal event because neural activity during seizures becomes more synchronous and predictable (Mammone et al. 2015; Nicolaou and Georgiou 2012; Tibdewal et al. 2017). This property of ictal discharges has been used extensively to detect and predict seizures from neurophysiological recordings (Li et al. 2015; Sharmila et al. 2018). Previously, Urigu en et al analysed background scalp EEG recordings in persons with idiopathic generalised epilepsy (Urigu en et al. 2017). They found that Shannon Spectral Entropy (SSE) tends to be *higher* in persons with idiopathic epilepsy as compared to healthy controls. However, they also found that SSE *decreases* with the advent of a seizure and approaches values close to those of healthy controls over a span of time, which may last up to 8 years. Our results are in partial agreement with Urigu en et al, since we observed a decrease in entropy in rodent models after seizure onset.

### Contending with the second law of thermodynamics

As per the laws of thermodynamics entropy of an isolated system always increases over time. However, under certain conditions this law can be violated wherein entropy can decrease locally, only if it is compensated by an increase in entropy elsewhere such that the total entropy does not decrease (Kostic 2020).

Epileptic seizures are marked by a paradoxical decrease in entropy, and the findings of this and other studies suggest that this decrease is sustained in the interictal states as well. However, the diverse aetiologies of epilepsy -such as genetic mutations, stroke, head injuries and infection – suggest a pre-seizure state characterized by transient or sustained higher entropy (as also reported by Urigu en et al). These entropy ‘spikes’ may be a manifestation of ionic imbalances, neuroinflammation, maladaptive plasticity, etc. Therefore, it is possible that the onset of seizures may represent an attempt to resolve these entropy ‘spikes’. This may help explain the convergence of diverse pathophysiological processes and insults into the common outcome state of a seizure.

### Implications for early life epilepsy

It is well established that an earlier age of seizure onset is predictive of greater risk of developing drug resistance, mortality (Berg et al. 2019; Nickels 2019) and is also associated with poor neurodevelopmental outcomes (Sto dberg et al. 2022) and intellectual disability (Vignoli et al. 2016). A pivotal study by Saxe et al calculated the brain entropy of resting state fMRI signals from a very large cohort of healthy individuals and found that higher entropy was associated with higher intelligence. They concluded that complex cognitive behaviour is strongly linked with access to varied neural states, and this information can be derived from the resting state signal alone (Saxe et al. 2018). Therefore, results from our study offer a possible explanation for the association between early life epilepsies and poor intellectual development. Indeed, we observed that early onset was associated with lower brain entropy which carried over well into later life. This adds greater impetus to timely diagnose and treat epilepsies with an early age of onset.

Given the retrospective nature of this study, we were unable to retrieve healthy control data for human MEG recordings, especially since a large fraction of the study cohort was paediatric, and therefore not apt for comparison with healthy adult subjects. However, comparison of sample entropy in individuals with epilepsy and healthy subjects can be a topic of future research in this area.

## Supporting information

eFigure1

## Acknowledgements

This research used TAURUS, the Aston University High Performance Computing facility. We would like to thank Peter Taylor and CNNP Lab, Newcastle University for greatly supporting this study. KK was supported by Institute of Health and Neurodevelopment, Aston University for completing this work. We would also like to thank the Aston Biomedical Facility for supporting the collection of rodent data. KK would like to thank Enrico Amico for a very insightful discussion on topics concerning this manuscript.

