## Supplementary material for "Entropy of the resting state cortex in epilepsy": eFigure1

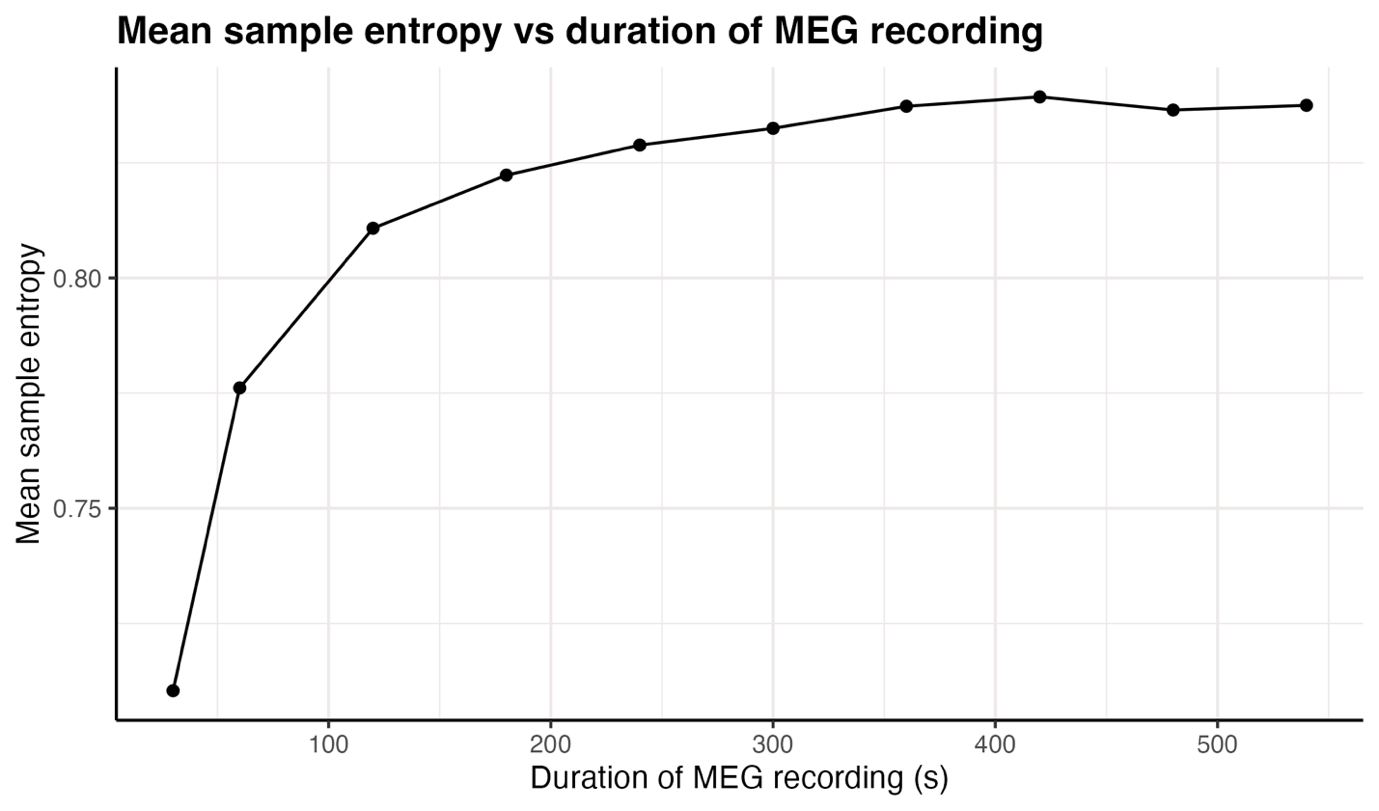

**eFigure1: Mean sample entropy across all patients varies in response to the duration of MEG source localized time series. The mean value of entropy stabilizes around 360s. In this study 540s (9 minutes) of time series were analysed.**
